# *Arabidopsis thaliana* subclass I ACTIN DEPOLYMERIZING FACTORs regulate nuclear organization and gene expression

**DOI:** 10.1101/2023.04.19.537409

**Authors:** Tomoko Matsumoto, Takumi Higaki, Hirotomo Takatsuka, Natsumaro Kutsuna, Yoshiyuki Ogata, Seiichiro Hasezawa, Masaaki Umeda, Noriko Inada

**Affiliations:** Graduate School of Agriculture, Osaka Metropolitan University, 1-1 Gakuen-cho, Naka-ku, Sakai, Osaka 599-8531, Japan; Graduate School of Science and Technology, Kumamoto University, Kumamoto 860-8555, Japan; International Research Organization for Advanced Science and Technology (IROAST), Kumamoto University, Kumamoto 860-8555, Japan; Graduate School of Science and Technology, Nara Institute of Science and Technology, Takayama-cho 8916-5 Ikoma, Nara 630-0192 Japan; LPIXEL, Inc., 1-6-1 Otemachi, Chiyoda-ku, Tokyo 100-0004, Japan; Graduate School of Science and Engineering, Hosei University, Tokyo 102-8160, Japan

**Author notes:** Author in correspondence. Present address:School of Biological Science and Technology, College of Science and Engineering, Kanazawa University, Kakuma-machi, Kanazawa 920-1192, Japan (Hirotomo Takatsuka).

## Abstract

ACTIN DEPOLYMERIZING FACTOR (ADF) is a conserved protein that regulates the organization and dynamics of actin microfilaments. Eleven ADFs in the *Arabidopsis thaliana* genome are grouped into four subclasses, and subclass I ADFs, ADF1–4, are all expressed throughout the plant. Previously, we showed that subclass I ADFs function in the regulation of the response against powdery mildew fungus as well as in the regulation of cell size and endoreplication. Here, we report a new role of subclass I ADFs in the regulation of nuclear organization and gene expression. Through a microscopic observation of epidermal cells in mature leaves, we found that the size of chromocenters in both *adf4* and transgenic lines where expression of subclass I *ADF*s are downregulated (*ADF1-4Ri*) was reduced compared with that of wild-type Col-0. *A. thaliana* possesses eight *ACTIN* genes, among which *ACT2*, *-7*, and *-8* are expressed in vegetative organs. The chromocenter size in *act7*, but not in the *act2/8* double mutant, was enlarged compared with that in Col-0. Microarray analysis revealed that 1,818 genes were differentially expressed in *adf4* and *ADF1-4Ri*. In particular, expression of 22 nucleotide-binding leucine-rich repeat (*NLR*) genes, which are involved in effector-triggered plant immunity, was reduced in *adf4* and *ADF1-4Ri*. qRT-PCR confirmed the altered expressions shown with microarray analysis. Overall, these results suggest that ADF regulates various aspects of plant physiology through its role in regulation of nuclear organization and gene expression. The mechanism how ADF and ACTIN regulate nuclear organization and gene expression is discussed.

## INTRODUCTION

ACTIN DEPOLYMERIZING FACTOR (ADF) is an ancient protein that is conserved among eukaryotes. ADF regulates both the organization and dynamics of actin microfilaments (AFs) by severing and depolymerizing them at the minus end (Inada, 2017). Analyses using various plants including *Arabidopsis thaliana* have revealed that ADFs function in regulating many aspects of plant physiology, including growth and development (Dong et al. 2001, Henty et al. 2011, Inada et al. 2021), flowering (Dong et al. 2001, Burgos-Rivera et al. 2008), immunity against bacterial (Tian et al. 2009, Porter et al. 2012) and fungal pathogens (Inada et al. 2016, Zhang et al. 2017, Sun et al. 2021), root-knot nematodes (Clément et al. 2009), and aphids (Mondal et al. 2018), rhizobial infection and nodule formation (Ortega-Ortega et al. 2020), and responses against drought (Qian et al. 2019, Sengupta et al. 2019), salt stress (Sengupta et al. 2019, Wang et al. 2021), and low temperature (Xu et al. 2021). However, it is unknown how ADFs regulate these broad aspects of plant physiology.

AFs mainly function underneath the plasma membrane by regulating intercellular transport (Henty-Ridilla et al. 2013); however, recent research using animal and insect cells has shown the roles of actin and actin-binding proteins in the nucleus and in the regulation of gene expression. The actin monomer has been shown to be a functional component of RNA polymerases and chromosome remodeling complex (Percipalle and Vartiainen 2019). Furthermore, recent development of nuclear AF visualization method revealed that nuclear AFs are transiently formed upon various stimuli, such as treatments with serum and reagents that induce DNA damage (Baarlink et al. 2013, Hurst et al. 2019) and contribute to the regulation of gene expression (Percipalle and Vartiainen 2019). Cytoplasmic AFs also regulate gene expression by associating with chromatin through nuclear envelope-embedded protein complexes and nuclear lamina (Davidson and Cadot 2021).

Interest has increased in plants in the role of actin and actin-binding proteins in the nucleus and in the regulation of gene expression (Porter and Day 2013, Porter and Day 2016, Inada 2017). Immunostaining using antibodies that label *A. thaliana* ACTIN2/8 (ACT2/8) and ACT7 showed accumulation of these actins within isolated nuclei (Kandasamy et al. 2010). Among the 11 members of ADFs in *A. thaliana* (Ruzicka et al. 2007, Inada, 2017), a knockout mutant of *ADF4* showed an increased susceptibility against the bacterial pathogen *Pseudomonas syringae* DC3000 harboring AvrPphB (Tian et al. 2009). This increased susceptibility is correlated with a decreased expression of Flg22-Induced Receptor Kinase 1 (*FRK1*) and Resistance to *Pseuromonas syringae* 5 (*RPS5*), which encode bacterial flagellin-induced receptor kinase and a resistance protein required for recognition of AvrPphB, respectively (Porter et al. 2012). The loss of *A. thaliana ADF9* expression causes reduced expression of *FLOWERING LOCUS C* (*FLC*), which is a master repressor of the transition from the vegetative to reproductive growth, and increased expression of *CONSTANS* (*CO*), *FLOWERING LOCUS T* (*FT*), *SUPRESSOR OF CONSTANS1* (*SOC1*), and *LEAFY* (*LFY*), resulting in early flowering (Burgos-Rivera et al. 2008).

We previously demonstrated that *A. thaliana* subclass I ADFs (ADF1-4), particularly ADF4, are involved in response against powdery mildew fungus (Inada et al. 2016) as well as in the regulation of endoreplication and plant size (Inada et al. 2021) and suggested that subclass I ADFs may have a function in the nucleus and in the regulation of gene expression. The *ADF4* knockout mutant showed strong resistance against the *A. thaliana*-adapted powdery mildew fungus *Golovinomyces orontii*. This resistance was further enhanced in transgenic plants where expression of all subclass I *ADF*s was suppressed (*ADF1-4Ri*). By visualizing AFs with expression of GFP-hTalin (Takemoto et al. 2003), we found that the density and level of bundling of AFs at the cell surface are not significantly altered in both *adf4* and *ADF1-4Ri*. Observation of *adf4* plants expressing ADF4-GFP revealed nuclear localization of ADF4 in addition to colocalization with AFs at the cell surface. Expression of ADF4-GFP complemented the *adf4 G. orontii*-resistant phenotype. However, expression of ADF4-GFP conjugated with nuclear exporting signal did not complement *adf4*, indicating that nuclear localization of ADF4 is important for response against *G. orontii* (Inada et al. 2016). We found that *adf4* and *ADF1-4Ri* had an increased plant size and an enhanced endoreplication (Inada et al. 2021). All known factors that regulate endoreplication are localized to the nucleus (discussed in Inada et al. 2021), further suggesting the role of subclass I ADFs in the nucleus or in the regulation of events occurring within the nucleus.

In this study, we characterized the nuclear morphology and gene expression profile of *A. thaliana adf4* and *ADF1-4Ri*. Nuclei in epidermal cells of *A. thaliana* mature leaves contain distinctive chromocenters that are strongly stained with DNA-binding fluorescent dyes (Fransz et al. 2002). By a close observation using confocal laser scanning microscope, we found that the size of these chromocenters of nuclei in the epidermal cells of mature leaves was significantly reduced in *adf4* and *ADF1-4Ri*. The *A. thaliana* genome contains eight ACTIN members, of which *ACTIN2*, −*7*, and −*8* are expressed in vegetative tissues (McKinney and Meagher 1998). We found that *act7*, but not *act2/8* mutants showed increased chromocenter size. Microarray analysis of wild-type Col-0, *adf4*, and *ADF1-4Ri* mature leaves showed that expression of a large number of genes was altered in both *adf4* and *ADF1-4Ri*. These results suggest that actin and ADF are involved in the regulation of nuclear organization and of gene expression.

## RESULTS

### The nuclear organization was altered in both *adf4* and *ADF1-4Ri*

For the first step to understand the role of ADFs in the nucleus or in the regulation of events occurring in the nucleus, we took a close observation of nuclei using a confocal laser scanning microscope. In fixed-*A. thaliana* mature leaves, 4ʹ,6-diamidino-2-phenylindole (DAPI)-stained nuclei contain well-defined chromocenters (Fransz et al. 2002). These chromocenters consist of heterochromatin regions including centromeric and pericentromeric domains as well as nucleolar organizing regions (NORs) that are composed of ribosomal DNA tandem repeats located in the short arms of chromosomes 2 and 4 (Fransz et al. 2002). NORs are located close to the nucleolus while centromeres are located near the nuclear periphery. By observing epidermal nuclei in mature leaves, we noticed that chromocenters in both *adf4* and *ADF1-4Ri* were smaller than those in Col-0 (Figs. 1A–C). No significant alteration in the shape and size of nuclei was noticed.

**Figure 1.**
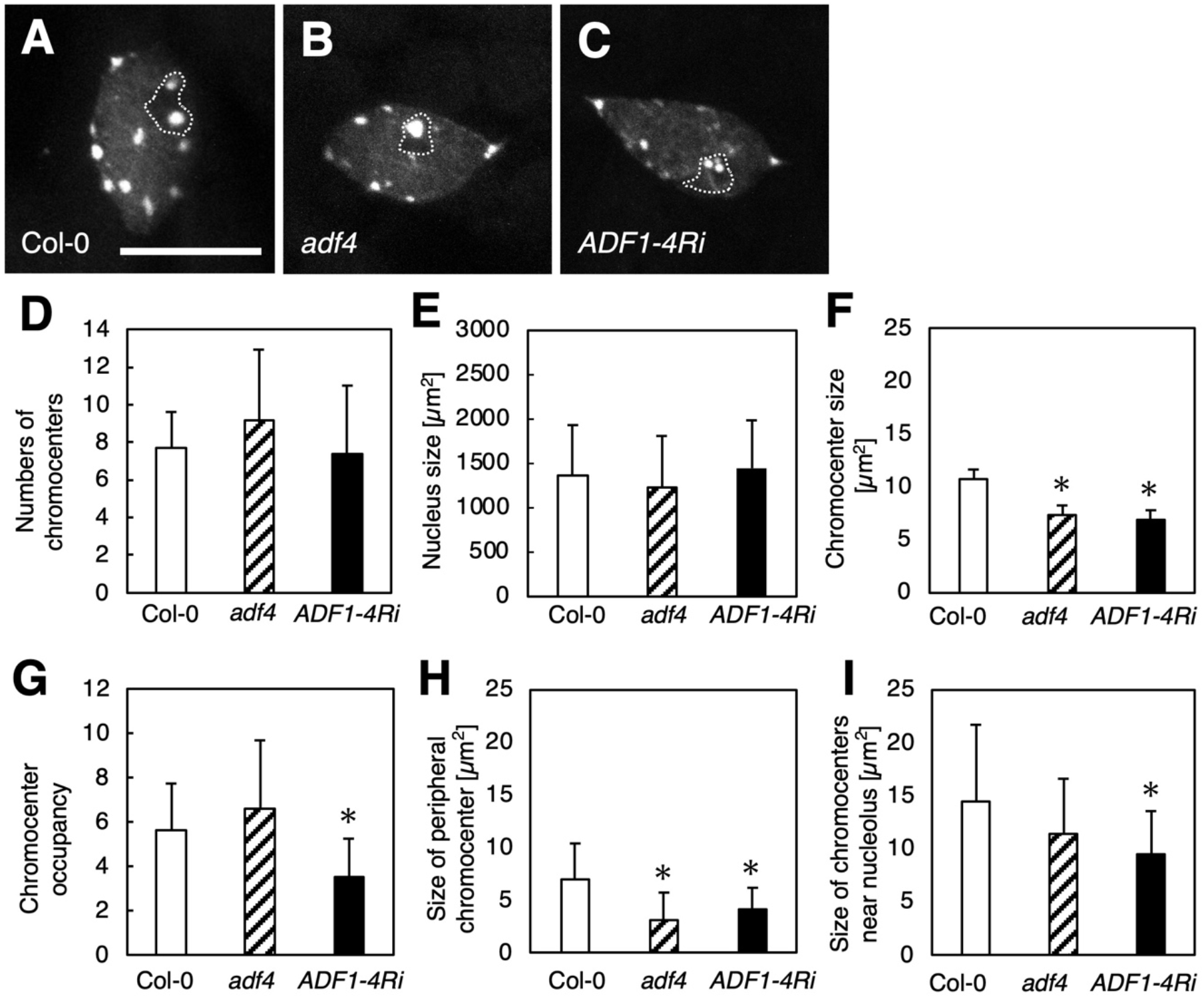
The size of chromocenters was reduced in *adf4* and *ADF1-4Ri* mature leaves. The fluorescence micrographs of DAPI-stained nuclei in epidermal cells of mature leaves in Col-0 (**A**), *adf4* (**B**), and *ADF1-4Ri* (**C**). Dotted lines indicate nucleoli. Bar indicates 5 µm. (**D**) Numbers of chromocenters, (**E**) the size of nuclei, (**F**) the size of chromocenters, (**G**) the percentage of chromocenter area in nuclei, (**H**) the size of chromocenters located at the nuclear periphery, and (**I**) the size of chromocenters located near the nucleoli in Col-0, *adf4*, and *ADF1-4Ri* were quantified using fluorescence micrographs. Twentyfour −27 nuclear images were used for quantitative analysis. Comparison with Col-0 was performed by Student’s *t*-test. Asterisks indicate that there is a significant difference (* *P*<0.05).

To quantify the size of chromocenters, maximum projection images produced from Z-stack DAPI-stained nuclei images were processed to evaluate the chromocenter regions. The number of chromocenters per nucleus (Fig. 1D), average size of nucleus (Fig. 1E), average size of chromocenters (Fig. 1F), and percentage of area occupied by chromocenters in the nucleus (Fig. 1G) were determined. Wild-type Col-0 nuclei contain 8–10 chromocenters as in previous observations (Fransz et al. 2002). As shown in Fig. 1D, the number of chromocenters per nucleus was unaltered in *adf4* and *ADF1-4Ri*. The size of nuclei in *adf4* and *ADF1-4Ri* was comparable with that in Col-0 (Fig. 1E). However, the average size of chromocenters in both *adf4* and *ADF1-4Ri* was significantly reduced compared with that in Col-0 (Fig. 1F). The percentage of chromocenter occupancy in the nucleus was also significantly reduced in *ADF1-4Ri* but not in *adf4* (Fig. 1G). In *ADF1-4Ri*, expression of all of four subclass I *ADF*s is suppressed by expression of RNAi construct targeting *ADF1* through *ADF4* (Tian et al. 2009). Four independent lines have been established for *ADF1-4Ri* (Tian et al. 2009), and all *ADF1-4Ri* lines showed significant reduction in chromocenter size (Fig. S1). µ

Close examination of the obtained images showed that the NOR chromocenters tended to be larger than those near the nuclear periphery in *adf4* and *ADF1-4Ri* (Figs. 1A–C; nucleoli are indicated by dotted lines). We examined the size of NOR chromocenters separately from chromocenters located at the nuclear periphery. The size of chromocenters at the nuclear periphery was significantly reduced in both *adf4* and *ADF1-4Ri* compared with the size of those in Col-0 (Fig. 1H). However, the NOR chromocenter size was not significantly altered in *adf4* compared with that in Col-0, whereas a significant reduction in chromocenter size was observed in *ADF1-4Ri* (Fig. 1I). This result suggests that chromocenters near the nucleoli are affected differently from those near the nuclear periphery in *adf4* and *ADF1-4Ri*.

Subclass I *ADF*s are more strongly expressed in mature leaves than in cotyledons (Ruzicka et al. 2007). We examined the chromocenter size of both cotyledons and mature leaves (first leaves) to understand the relationship between loss of subclass I *ADF* expression and chromocenter size reduction. The chromocenter size in the cotyledons of *adf4* and *ADF1-4Ri* was comparable to that in wild-type Col-0 (Figs. 2A–D), whereas the size of chromocenters in the first leaves of *ADF1-4Ri* was significantly reduced (Figs. 2E–H). qRT-PCR confirmed the reduction in expression of all of subclass I *ADF*s in the first leaves (Fig. S2). On the other hand, the level of expression suppression in cotyledons varies among members of subclass I *ADF*s. While *ADF1* and *ADF2*, which showed higher expression in cotyledons compared to first leaves in our growth condition, were suppressed in *ADF1-4Ri* as in first leaves (Figs. S2A and S2B), the expression of *ADF3* and *ADF4*, which was very low in Col-0 cotyledons, was increased in *ADF1-4Ri* cotyledons (Figs. S2C and S2D). Those results may suggest that members of subclass I *ADF*s differently contribute to regulation of chromocenter size.

**Figure 2.**
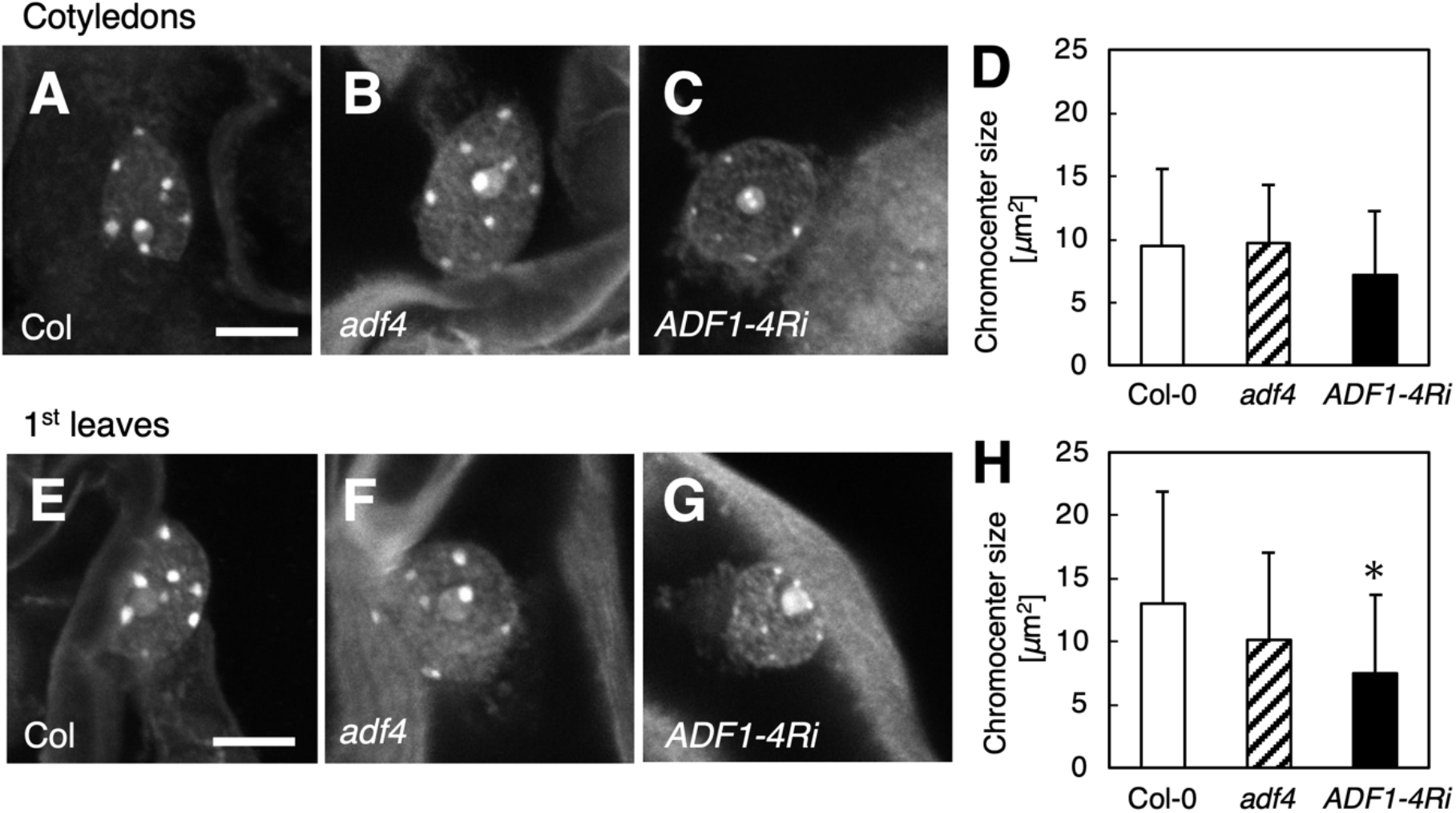
The size of chromocenters was less affected in cotyledons of *adf4* and *ADF1-4Ri*. The fluorescence micrographs of DAPI-stained nuclei in epidermal cells of cotyledons in Col-0 (**A**), *adf4* (**B**), and *ADF1-4Ri* (**C**). (**D**) The size of chromocenters in Col-0, *adf4*, and *ADF1-4Ri* cotyledons. Twelve nuclear images each were used to obtain average size of chromocenters. The fluorescence micrographs of DAPI-stained nuclei in epidermal cells of 1^st^ leaves in Col-0 (**E**), *adf4* (**F**), and *ADF1-4Ri* (**G**). (**H**) The size of chromocenters in Col-0, *adf4*, and *ADF1-4Ri* 1^st^ leaves. Twelve nuclear images each were used to obtain average size of chromocenters. Bars indicate 5 µm. Comparison with Col-0 was performed by Student’s *t*-test. Asterisks indicate that there is a significant difference (* *P*<0.05).

The reduction in chromocenter size could be caused by the decondensation of chromocenters. We performed in situ fluorescence hybridization (FISH) using a probe specifically labeling centromeric repetitive DNA to examine this possibility. Although most FISH labeling showed a condensed pattern, several nuclei showed diffused FISH labeling (Fig. S3), suggesting that the diffusion of heterochromatin caused a reduction in chromocenter size in *adf4* and *ADF1-4Ri*.

We also noticed that a portion of the nuclei in *ADF1-4Ri* contained chromocenters that tended to be clustered compared with the relatively random distribution in Col-0 nuclei (Figs. S4A and S4B). We quantified this chromocenter organization by measuring the distance of the two chromocenters (Fig. S4C). The median value of the distances between any closest chromocenter of nuclei was lower in *ADF1-4Ri* than that in Col-0. As the number of chromocenters per nucleus was unaltered in *ADF1-4Ri,* the positioning of chromocenters of *ADF1-4Ri* was disturbed and more aggregated than that in Col-0.

Thus, nuclear organization was altered in both *adf4* and *ADF1-4Ri* but more significantly in *ADF1-4Ri*.

### The size of chromocenters was altered in *act7* but not in *act2/8*

Among the 10 *ACT* genes in the *A. thaliana* genome (McDowell et al. 1996a), *ACT2*, *-7*, and *-8* are expressed in the vegetative tissues (An et al. 1996, McDowell et al. 1996b), where subclass I *ADF*s are mainly expressed (Ruzicka et al. 2007). ACT2 and ACT8 are highly similar in amino acid sequences and are functionally equivalent (Kandasamy et al. 2009). We reasoned that if ADFs regulate nuclear organization via AFs, it is possible that *act* mutants also show alterations in nuclear organization. We therefore examined the nuclear morphology of the *act2-1*/*act8-2* double mutant (*act2/8*) as well as that of *act7-4* single mutant (*act7*)(Figs. 3A–C). No significant alterations were observed in the number of chromocenters per nucleus (Fig. 3D) or in the nuclear size (Fig. 3E) in *act2*/*8* or *act7*. We noticed that the size of chromocenters tended to be larger in *act7* (Fig. 3C) than that in Col-0 (Fig. 3A). However, quantitative analysis of chromocenter size showed no significant difference among Col-0, *act2/8*, and *act7* when all chromocenters were included for analysis (Fig. 3F). The percentage of area occupied by chromocenters in the nucleus was also unaltered in *act2/8* and *act7* (Fig. 3G). When chromocenters at the nuclear periphery and those near the nucleolus were separately analyzed, the chromocenters at the nuclear periphery were significantly larger in *act7* than those in Col-0 (Fig. 3H), whereas the chromocenters near the nucleolus were comparable among Col-0, *act2/8*, and *act7* (Fig. 3I).These results further confirmed that actin disturbance affects the size of chromocenters, and that chromocenters near the nucleolus were regulated differently from those at the nuclear periphery.

**Figure 3.**
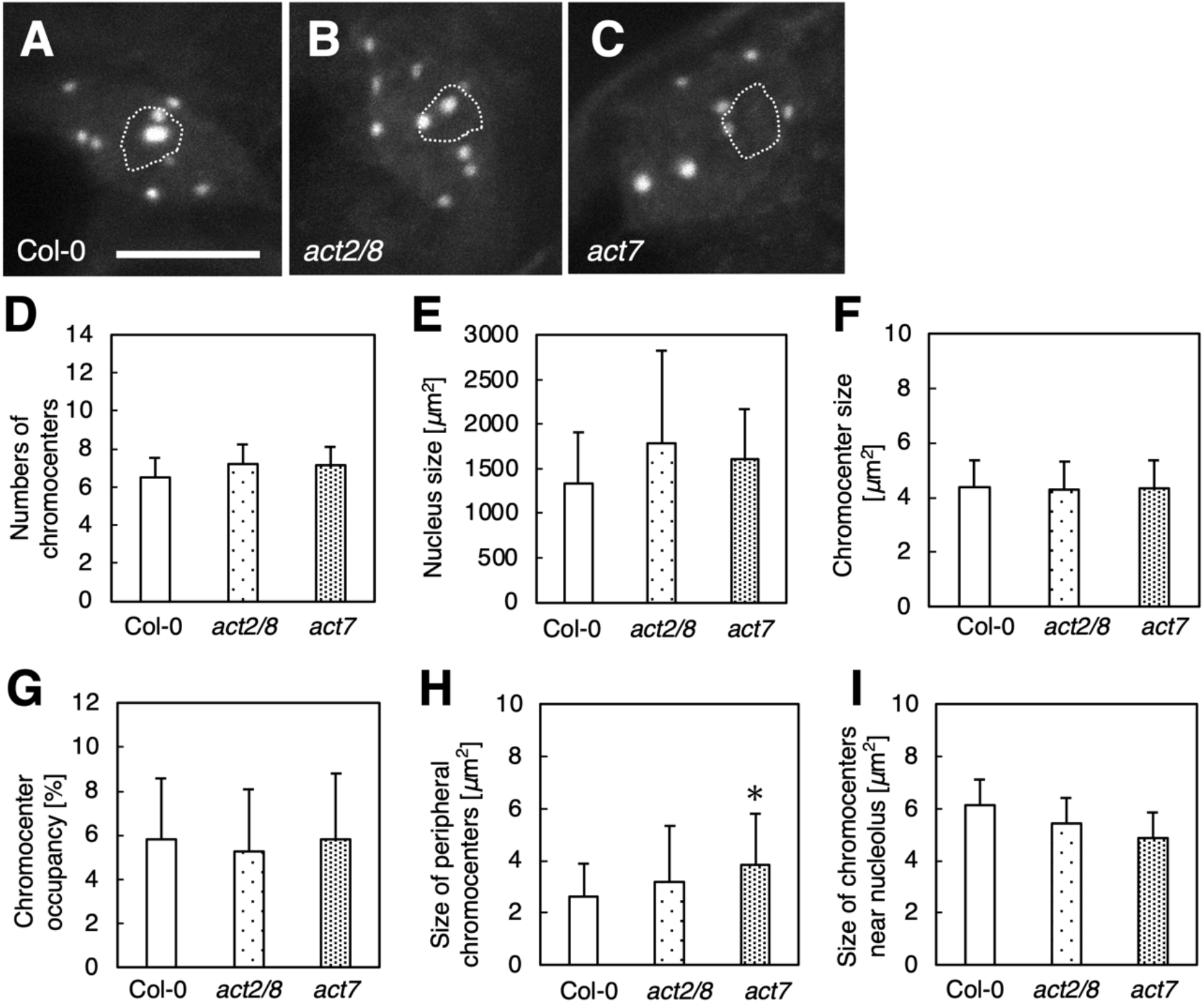
The size of chromocenters was increased in *act7* mature leaves. The fluorescence micrographs of DAPI-stained nuclei in epidermal cells of mature leaves in Col-0 (**A**), *act2/8* (**B**), and *act7* (**C**). Dotted lines indicate nucleoli. Bar indicates 5 µm. (**D**) Numbers of chromocenters, (**E**) the size of nuclei, (**F**) the size of chromocenters, (**G**) the percentage of chromocenter area in nuclei, (**H**) the size of chromocenters located at the nuclear periphery, and (**I**) the size of chromocenters located near the nucleoli in Col-0, *act2/8*, and *act7* were quantified using fluorescence micrographs. Sixteen – 19 nuclear images were used for quantitative analysis. Comparison with Col-0 was performed by Student’s *t*-test. Asterisks indicate that there is a significant difference (* *P*<0.05).

We then analyzed chromocenters in *fiz1*, dominant negative mutant of *ACT8*. In *fiz1* homozygous mutant, the organization of AFs is significantly disrupted, and AFs became shorter and thicker comparing to wild type or *fiz1* heterozygous mutant (Kato et al. 2010). As shown in Fig. S5, there was no difference in number of chromocenters per nucleus, nuclear size, and chromocenter size between wild type Col-0 and *fiz1* homozygous mutant.

### Gene expression profile was significantly altered in both *adf4* and *ADF1-4Ri*

The altered nuclear organization in *adf4* and *ADF1-4Ri* described above may affect gene expression. To test this idea, we performed a large-scale gene expression analysis using the Arabidopsis Gene 1.0 ST array. RNA extracted from mature leaves of 4-week-old Col-0, *adf4*, and *ADF1-4Ri* was used for microarray analysis. Two independent sets of plants were used for microarray analysis for biological replication. Genes that showed more than two-fold differences in expression levels in both data set were extracted. With this criterion, we found that expression of 911 genes in total was upregulated in *adf4* and *ADF1-4Ri*, and that of 907 genes in total was downregulated in *adf4* and *ADF1-4Ri* (Figs. 4A and 4B). Among these, expression of 137 genes was upregulated in both *adf4* and *ADF1-4Ri*, whereas that of 232 genes was downregulated in both *adf4* and *ADF1-4Ri* (Tables S1 and S2).

**Figure 4.**
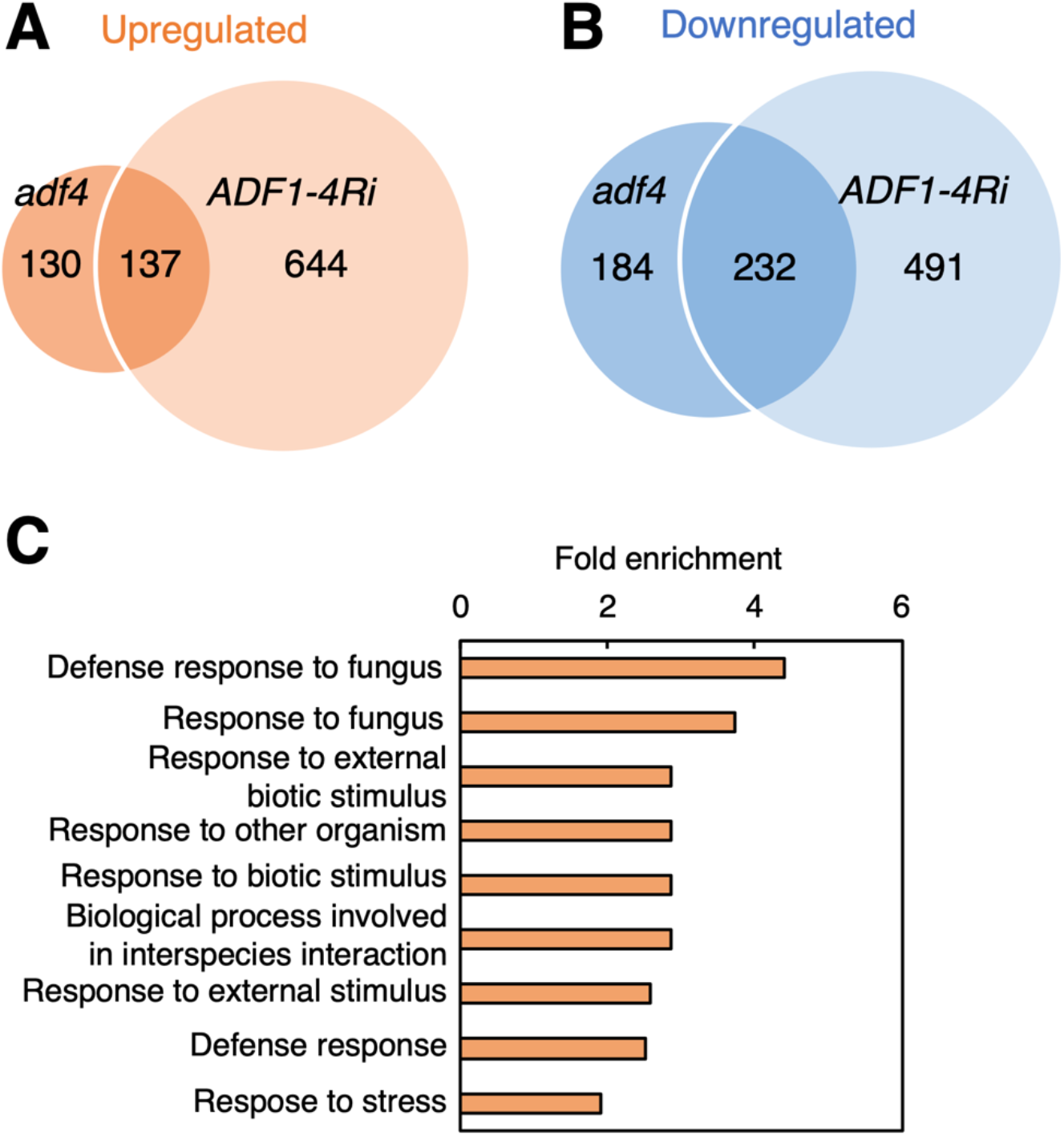
Microarray analysis showed altered expression of many genes in *adf4* and *ADF1-4Ri* mature leaves. Venn diagrams of upregulated (**A**) and downregulated (**B**) genes in *adf4* and *ADF1-4Ri*. (**C**) GO analysis for genes that showed altered expression in both *adf4* and *ADF1-4Ri* (genes including both upregulated and downregulated genes).

Gene Ontology analysis performed for these up- or downregulated genes in both *adf4* and *ADF1-4Ri* revealed enrichment of biotic-responsive genes (Fig. 4C). We also found that downregulated genes included many nucleotide-binding-leucine-rich repeat (*NLR*) genes (Table S3). NLRs function in the recognition of pathogen effectors that are delivered into plant cells. Recognition of effectors by NLRs triggers plant immunity (Pok Man Ngou et al. 2022). Misregulation of *NLR* expression can cause autoimmunity and growth defects, and the expression level of *NLR* genes is therefore maintained at a low level in uninfected plants (Fick et al. 2022). We found that the expression of 22 *NLR* genes were further downregulated in both *adf4* and *ADF1-4Ri* (Table S3). The list of downregulated *NLR*s in both *adf4* and *ADF1-4Ri* included *RPS5* (At1g12220), whose expression is reported to be downregulated in *adf4* (Porter et al. 2012). The expression of five *NLR*s were downregulated in *adf4* but not in *ADF1-4Ri*. Twenty-one *NLR*s showed reduced expression in *ADF1-4Ri* but not in *adf4* (Table S3). Upregulated *NLRs* included one *NLR* in *adf4*, and eight *NLR*s in *ADF1-4Ri*. No *NLRs* were upregulated in both *adf4* and *ADF1-4Ri* (Table S3). This result suggests that each member of subclass I ADFs contributes differently to the regulation of expression of different *NLR*s.

### qRT-PCR confirmed the altered gene expression in *adf4* and *ADF1-4Ri*

As we only took biological duplications for microarray analysis, we verified the microarray data by performing qRT-PCR for genes that are up- or downregulated in both *adf4* and *ADF1-4Ri* by using *ACT8* as an internal control. We selected genes for qRT-PCR analysis based on three criteria: genes whose functions have been reported; genes where the expression level is sufficiently high in Col-0 (based on eFP browser analysis; http://bar.utoronto.ca/efp2/Arabidopsis/Arabidopsis_eFPBrowser2.html), and genes that possess specific sequences that have low homology with other genes.

For upregulated genes, At1g13210, At1g18710, At1g45145, At1g52000, At1g52400, and At3g45860 were selected. At1g13210 encodes autoinhibited Ca^2+^/ATPase II (ACA.l), which functions as a P4-type ATPase that maintains asymmetry between the intra- and extracellular lipid layers in phospholipid membrane bilayers (Axelsen et al. 2001). At1g18710 encodes MYB DOMAIN PROTEIN 47 (MYB47), a member of R2R3-MYB proteins that function in plant primary and secondary metabolism, cell fate determination, development, and responses to both biotic and abiotic stresses (Dubos et al. 2010). *MYB47* expression is reported to be induced by wounding and methyl jasmonate (MeJA) treatment (Devoto et al. 2005). At1g45145 encodes thioredoxin *h5* (TRX5), a member of *h*-type thioredoxin. *A. thaliana TRX5* is required for the response to victorin, an effector produced by a fungus *Cochliobolus victoriae*, which causes Victoria blight of oats (Sweat and Wolpert. 2007). At1g52000, encoding a member of jacalin-related lectin family (Eggermont et al. 2017), is reportedly upregulated by treatment with both jasmonic acid and MeJA (Sasaki-Sekimoto et al., 2005) as well as by inoculation with fungal pathogen *Fusarium oxysporum* (Zhu et al. 2013). At1g52400 encodes BETA GLUCOSIDASE 18 (BGLU18), a protein that localizes the endoplasmic reticulum (ER) and accumulates in ER bodies (Ogasawara et al. 2009). BGLU18 is also known to mediate hydrolysis of abscisic acid (ABA)-glucose ester to produce active ABA (Han et al. 2020). At3g45860 encodes CYSTEINE-RICH RECEPTOR-LIKE PROTEIN KINASE 4 (CRK4). *CRK4* expression is induced by salicylic acid treatment and *P. syringae* pv. tomato DC3000 (*Pst*) infection (Chen et al. 2004). Overexpression of *CRK4* induces hypersensitive cell death (Chen et al. 2004) and confers resistance to the virulent bacteria *Pst* (Yeh et al. 2015). *CRK4* overexpression can enhance drought tolerance (Lu et al. 2016). The upregulation of expression of these genes in *adf4* and *ADF1-4Ri* was confirmed with qRT-PCR (Figs. 5A–F).

**Figure 5.**
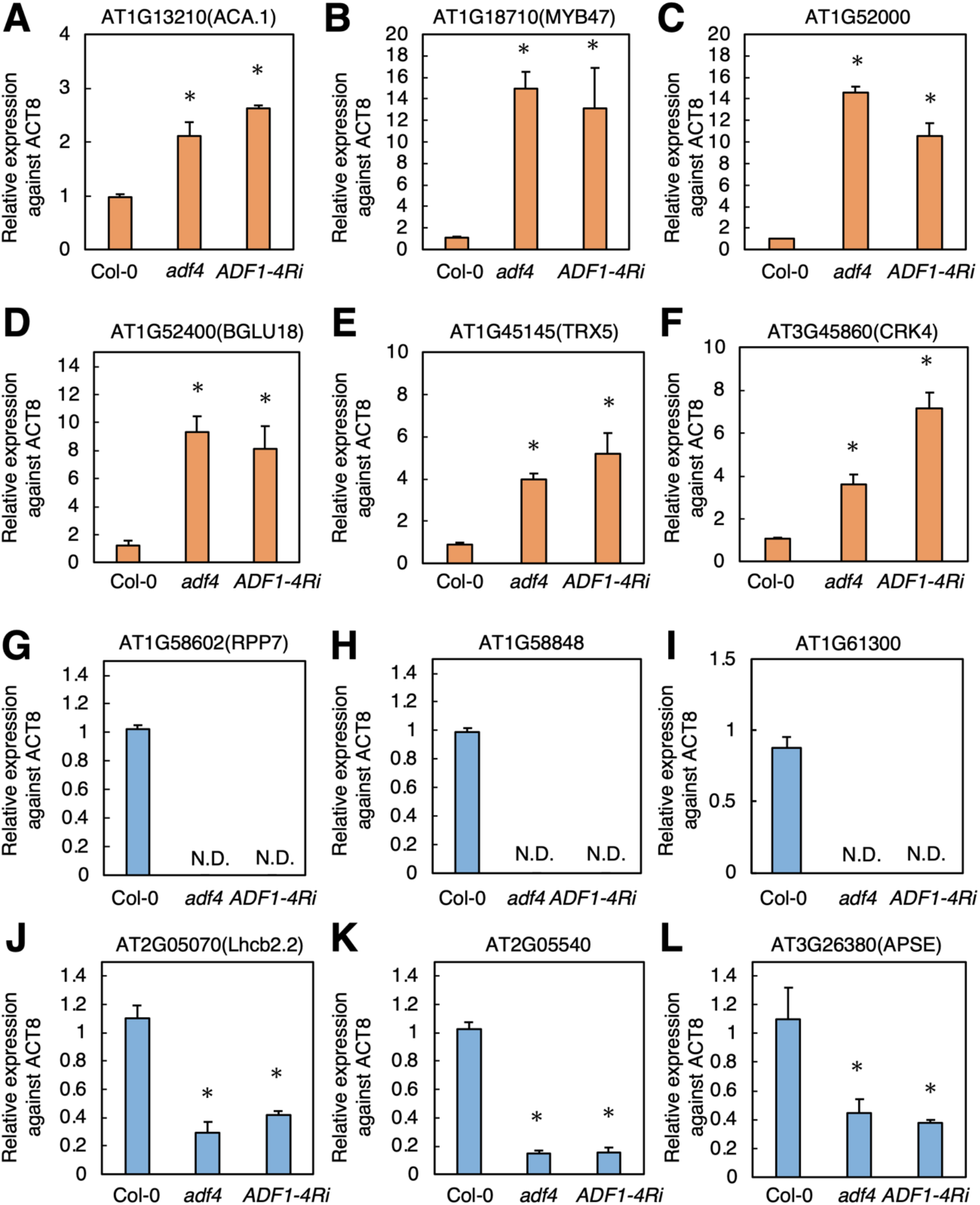
qRT-PCR confirmed alteration in gene expression in *adf4* and *ADF1-4Ri* shown by microarray analysis. The expression of At1g13210 (**A**), At1g18710 (**B**), At1g52000 (**C**), At1g52400 (**D**), At1g45145 (**E**), and At3g45860 (**F**) was upregulated in both *adf4* and *ADF1-4Ri*. The expression of At1g58602 (**G**), At1g58848 (**H**), At1g61300 (**I**), At2g05070 (**J**), At2g05540(**K**), and At3g26380 (**L**) was downregulated in both *adf4* and *ADF1-4Ri*. Standard deviation of experimental triplicates is shown. Comparison with Col-0 was performed by Student’s *t*-test. Asterisks indicate that there is a significant difference (* *P*<0.05). Representative data of biological triplicates are shown.

For downregulated genes, At1g58602, At1g58848, At1g61300, At2g05070, At2g05540, and At3g26380 were examined for their expression with qRT-PCR. At1g58602, At1g58848, and At1g61300 encode NLRs. NLRs can be divided into three groups according to their N-terminal domains, namely, those with Toll/interleukin-1 receptor (TIR) domain, coiled-coil (CC) domain (CNL), and RESISTANCE TO POWDERY MILDEW (RPW) 8-like CC domain (Pok Man Ngou et al. 2022). At1g58602, At1g58848, and At1g61300 are all CNL. At1g58602 encodes RPP7 (for resistance to *Peronospora paracitica* 7), which is required for resistance against oomycete *Hyaloperonospora paracitica* Hiks1 (McDowell et al. 2000). The expression of At1g58602, At1g58848, and At1g61300 in wild-type Col-0 was low but detectable (Figs. 5G–I). However, in *adf4* and *ADF1-4Ri*, the expression of these *NLRs* was reduced to an undetectable level (Figs. 5G–I), confirming the microarray results.

At2g05070 encodes light-harvesting chlorophyll-binding 2.2 (Lhcb2.2). *Lhcb1*, *Lhcb2*, and *Lhcb3* genes encode polypeptides for trimeric LHCII (Jansson 1999). Lhcb1 is the most abundant among these Lhcb proteins, and Lhcb2 plays a critical role in state transitions (Pietrzykowska et al. 2014). Three genes in *A. thaliana* Col-0 encode *Lhcb2*, namely, *Lhcb2.1* (At2g05100), *Lhcb2.2* (At2g05070), and *Lhcb2.3* (At3g27690). As *Lhcb2.1*, *2.2*, and *2.3* encode identical proteins (Jansson 1999), we designed primers that detect the 3’-UTR of *Lhcb2.2* that showed a decrease in *Lhcb2.2* expression in both *adf4* and *ADF1-4Ri* (Fig. 5J). The expression of *Lhcb2.1* (At2g05100) and *Lhcb2.3* (At3g27690) also exhibited a degree of decrease in expression according to microarray analysis (Table S4). At2g05540 encodes a member of glycine-rich protein family, although the function is largely unknown. Expression of At2g05540 is regulated by SWI/SNF chromatin remodeling protein AtCHR12 (Mlynárová et al. 2007) and by the SET domain protein, SDG4, which maintains the level of histone H3 lysine 4 (H3K4) and H3K36 methylation (Cartagena et al. 2008). At3g26380 encodes β-l-ARAPASE (APSE) that functions in the hydrolysis of β-l-arabinopyranosyl residues in arabinogalactan-proteins. A mutation in APSE causes inhibition of the elongation growth of hypocotyls (Imaizumi et al. 2017). The expression of both At2g05540 and At3g26380 was reduced in *adf4* and *ADF1-4Ri* (Figs. 5K and 5L). qRT-PCR using *UBQ11* as an internal control confirmed changes in expression of At1g18710, At3g45860, At1g58602 and At2g05070 in *adf4* and *ADF1-4Ri* (Fig.S6).

Overall, qRT-PCR analyses confirmed the microarray results, and we concluded that expression of many genes, including those encoding *NLR*s, was significantly altered in *adf4* and *ADF1-4Ri*.

## DISCUSSION

Herein, we showed that chromocenter organization was significantly altered in both *adf4* and *ADF1-4Ri*. The size of DAPI-stained chromocenters was significantly reduced in both *adf4* and *ADF1-4Ri*, and part of nuclei in *ADF1-4Ri* showed clustered chromocenters. This reduction in size was likely caused by the decondensation of chromocenters. In the *act7* mutant but not the *act2/8* mutant, the size of chromocenters was increased compared with that of Col-0. Microarray analyses revealed alterations in the expression of many genes in *adf4* and *ADF1-4Ri*. Overall, our results indicate that subclass I ADFs and ACT function in the regulation of nuclear organization and gene expression.

Based on previous research using mammalian cells, we propose three possible mechanisms of ADF and ACT regulating nuclear organization and gene expression. First possibility is that ACT monomer in the nucleus regulate chromatin organization and gene expression. In mouse embryonic fibroblasts, loss of β-actin showed an increase in nuclear area and a decrease in DAPI fluorescence intensity in nuclei (Xie et al. 2018). Both immunofluorescence microscopy and western blotting showed that the amount of histone H3 trimethylated at lysine 9 (H3K9me3), a marker of facultative heterochromatin, was increased in β-actin knockout cells. Heterochromatin protein (HP)1α is a central component of heterochromatin in mammalian and *Drosophila* cells (Saksouk et al. 2015). The number of HP1α-labeled spot areas increased in β-actin knockout cells (Xie et al. 2018). This increase in the heterochromatin area in β-actin knockout cells is similar to the increased chromocenter size in *act7*. In mammalian cells, β-actin is an essential component of the chromatin remodeling complex BRG1/BRM-associated factor (BAF, mammalian SWI/SNF) (Kapoor and Shen 2014). β-actin is required for maximum ATPase activity of Brg1, an ATPase subunit of the BAF complex. Loss of β-actin significantly decreased DNA binding of Brg1 (Xie et al. 2018). Therefore, β-actin regulates heterochromatin organization through its DNA-binding activity of the BAF complex (Xie et al. 2018). In plants, orthologs of SWI/SNF subunits have been identified, and their functions have been characterized (Reyes 2014, Nishioka et al. 2020). The interaction between actin and BRM has not yet been reported in plants, but our results suggest that plant actin also plays a role in regulating the activities of the chromatin remodeling complex, which regulates heterochromatin organization and gene expression. In this scenario, loss of ADF stabilizes AFs, decreases the available actin monomers, thus affecting actin-mediating regulation of chromatin organization and gene expression.

Another possibility is that ADF and ACT regulate chromatin organization through regulation of cytoplasmic AFs, which are connected with chromatin through the nuclear envelope-embedded protein complex and nuclear lamina. Mammalian genomes have three ADFs, namely, Cofilin-1 (COF), ADF, and COF-2. The depletion of both COF and ADF expression induces severe deformation of nuclear shape in mice (Kanellos et al. 2015) and in several mammalian cultured cells, including HeLa cells (Wiggan et al. 2017). COF/ADF-depleted HeLa cells exhibit a significant reduction in the level of H3K27me3, which marks facultative heterochromatin at the nuclear periphery (Wiggan et al. 2017). The nuclear deformation in COF/ADF-silenced cells is associated with excessive activation of myosin-II in the cytoplasm and disruption of the nuclear lamina (Wiggan et al. 2017). The cytoplasmic AFs are connected with the nuclear envelope-embedded linker of the nucleoskeleton and cytoskeleton complex, which is composed of outer nuclear envelope-localized Klarsicht, ANC-1, and SYNE homology (KASH) domain-containing nesprins and inner nuclear envelope-localized SUN (Sad1 and UNC-84) domain-containing proteins (Tamura et al. 2015). SUN proteins interact with nuclear lamina (Tamura et al. 2015), and nuclear lamina interact with heterochromatin (Hoskins et al. 2021). Nuclei in COF/ADF-depleted mammalian cells show the disorganization of the nuclear lamina, and the authors concluded that excessive tension in the cytoplasm caused by the depletion of COF/ADF results in alterations in nuclear organization and gene expression (Wiggan et al. 2017).

The reduction of H3K27me3 level in COF/ADF-depleted HeLa cells resembles the reduction in the size of chromocenters in *adf4* and *ADF1-4Ri*. Our results that chromocenters at the nuclear periphery were more strongly affected compared with those near the nucleoli in *adf4*, *ADF1-4Ri* and *act7* (see Figs. 1H, 1I and 3H) also support this hypothesis that connection between cytoplasmic AFs and chromatin through nuclear envelope-embedded protein complex is important for regulation of chromatin organization. Indeed, the clustered chromocenters in *ADF1-4Ri* (Fig. S4) were similar to the polarized organization of chromocenters observed in *A. thaliana* mutants of KASH and SUN (Sakamoto et al. 2022). Our result of *fiz1* (Fig. S5), however, does not support this hypothesis. While cytoplasmic AFs are highly disrupted (Kato et al. 2010), *fiz1* did not show alteration in chromocenter size.

The result of *fiz1* supports the third hypothesis, in which the function of ADF in the nucleus, rather than in the cytoplasm, could contribute to the regulation of chromocenter organization. In mammalian cells, it has been shown that nuclear AFs are transiently formed upon various stimuli and regulate gene expression, DNA damage repair, and regulation of nuclear size (Percipalle and Vartiainen 2019). Subclass I ADFs and ACT may regulate heterochromatin organization and gene expression through a regulation of nuclear AFs, which has not been reported in plant cells yet. It is also possible that ADF and ACT function in a separate pathway to regulate chromatin organization and gene expression.

We showed that genes related to plant defense response are both up- and downregulated in *adf4* and *ADF1-4Ri*. Furthermore, we found that expression of many *NLR* genes are downregulated in *adf4* and *ADF1-4Ri*. Twenty-two *NLRs* showed reduced expression in both *adf4* and *ADF1-4Ri*. While our microarray analysis used uninfected mature leaves, it has been shown that *RPS5* expression, of which reduction was confirmed in our microarray, was reduced both in uninfected and *Pst* AvrPphB-infected *adf4* (Porter et al. 2012), suggesting that *NLR* expression is also affected in *adf4* and *ADF1-4Ri* during pathogen infection. Our result that expression of some *NLRs* was suppressed only in *adf4* or in *ADF1-4Ri* suggests that different ADF members affect expression of different sets of *NLR*s. This could explain previous results where loss of *ADF*s results in both increase or decrease in resistance depending on pathogens and that different ADF members affect pathogen response differently. Tian et al. reported that *A. thaliana adf4* showed an increased susceptibility against the bacterial pathogen *Pst* harboring AvrPphB, whereas *adf4* showed comparable susceptibility against *Pst* or *Pst* AvrB. Neither *adf1* nor *adf3* showed altered susceptibility against these pathogens. The susceptibility of *ADF1-4Ri* against *Pst* AvrPphB was also comparable with that of Col-0 (Tian et al. 2009). However, both *adf4* and *ADF1-4Ri* showed an increased resistance to *G. orontii* (Inada et al. 2016). In wheat (*Triticum aestivum* L.), the suppression of different members of ADFs has different effects on immunity against fungus causing stripe rust. Wheat genotype Suwon 11 is resistance to *Puccinia striiformis* f.sp. *tritici* CYR23 but is susceptible to *P. striiformis* f.sp. *tritici* CYR31. Knockdown of *TaADF7* (Fu et al. 2014) and *TaADF4* (Zhang et al. 2017) both increased a susceptibility against CYR23. By contrast, knockdown of *TaADF3* expression caused increased resistance against CYR31 (Tang et al. 2016). It is possible that TaADF3, 4, and 7 affect the expression of different set of *NLR*s, which results in the above-mentioned phenotypic differences.

Taken together, our results strongly suggests that *A. thaliana* subclass I ADFs regulate various aspects of plant physiology including pathogen response through their roles in regulation of nuclear organization and gene expression. Future research elucidating the mechanism how ADFs and actin regulate nuclear organization and gene expression will contribute to understand the new roles of plant cytoskeletons as well as new mechanisms regulating gene expression.

## MATERIALS AND METHODS

### Plant materials and growth condition

*adf4* (Garlic_823_A11b.1b.Lb3Fa) and *ADF1-4Ri* lines were described previously (Tian et al. 2009). The loss of *ADF4* transcript in *adf4* was previously examined (Tian et al. 2009). *act2-1/act8-2* double mutant and *act7-4* single mutant were described previously (Kandasamy et al. 2009). *act2-1* and *act7-4* were identified in mutant screenings of T-DNA insertion lines ((McKinney et al. 1995, Gilliland et al. 2003). *ACT2* transcript in *act2-1* is severely knocked down with <4% of wild type, while both *act7-4* and *act8-2* (GABi-Kat line 480C07) show low level of *ACT7* and *ACT8* transcripts, respectively, at 10∼15% of wild type (Kandasamy et al. 2009). *Arabidopsis* seeds were suspended in autoclaved 0.1% agarose and incubated at 4°C for vernalization for more than 1 day (up to 2 weeks) before direct sowing on 1:3 metromix:vermiculite in plastic pots. Entire pots were covered with plastic wrap for 1 week after sowing to maintain humidity and to encourage germination. Plants were grown at 22°C in a growth chamber under a 16-h light:8-h dark photoperiod (LH-411S, NK Systems, Osaka, Japan).

### Observation of chromocenters

For observation of chromocenters in epidermal cells, 4 week old mature leaves were fixed with 3:1 ethanol:acetic acid at R.T. overnight. Fixed leaves were then rinsed with milliQ three times before stained with 1 µg/mL DAPI. Images of DAPI-stained nuclei in epidermal cells were excited with 405 nm laser, and fluorescence between 410-585 nm was captured using confocal laser scanning microscope (Zeiss LSM700) that was equipped with ×63 objective lens (N.A. 1.4). Z-stack images were obtained with 0.5 μm step size.

### Quantitative image analysis of chromocenters

Obtained Z-stack confocal images of DAPI were combined using maximum intensity projection. To measure the chromocenter number and size, a binary image was obtained from the maximum intensity projection image by intensity thresholding. Before binarization, a Gaussian filter (sigma = 2 pixels) was applied to the images in order to improve the signal to noise ratio. Intensity thresholding was performed based on Yen’s method (Yen et al. 1995), and then the binary images were masked by manually segmented nuclear regions. Number and size of the chromocenter blobs in the binary images were measured. All image processing and analysis were performed by ImageJ software (Schneider et al. 2012).

### Microarray analysis

4-week old mature leaves were harvested and frozen in liquid N_2_. Total RNA was extracted using TRIzol reagent (Life Technologies Japan, Tokyo, Japan) and purified with an RNeasy Micro kit (Qiagen) following the manufacturers’ instructions. Arabidopsis Gene 1.0 ST arrays (Thermo Fisher Scientific) was used for microarray analysis. Quality of RNA was checked using RNA 600 Nano LabChip Kit (Agilent) following the manufacturer’s protocol. Sense-strand DNA target preparation was performed with WT Expression Kit (Ambion) and GeneChip WT Terminal labeling Kit and Controls Kit (Thermo Fisher Scientific) following the manufacturer’s protocols. Hybridization, washing, and staining procedures were performed on Fluidics Station 450 (Affymetrix) with GeneChip Hybridization, Wash, and Stain Kit (Thermo Fisher Scientific). GeneChips were scanned with GeneChip Scanner 3000 7G. Gene-level RMA sketch normalization was performed using standard settings for GeneChip Gene 1.0 ST arrays on Expression Console Version 1.3 (Affymetrix). Microarray data were quantified using Transcriptome Analysis Console 4.0 (Thermo Fisher Scientific). Expression levels were compared between Col-0 and *adf4* and between Col-0 and *ADF1-4Ri* by calculating average log2 ratios. Genes with significantly altered expression were defined as those with a fold change of ≥2 or ≤-2 from Col-0 for both *adf4* and *ADF1-4Ri*. GO analysis was performed using PANTHER 17.0 (Released 20220223). Microarray data were deposited in Gene Expression Omnibus (GEO) of National Center for Biotechnology Information (NCBI) and are accessible through GEO accession number GSE228396.

### qRT-PCR

Mature leaves were harvested from 4-week-old plants and frozen in liquid N_2_. Total RNA was extracted using TRIzol reagent (Life Technologies Japan, Tokyo, Japan) in accordance with the manufacturer’s protocol. cDNA was synthesized with the ReverTra Ace qPCR RT Master Mix with gDNA Remover (TOYOBO) using 100 ng of total RNA. qRT-PCR was carried out using THUNDERBIRD SYBR qPCR Mix (TOYOBO) on 7300 Real Time PCR System (Applied Biosystems). *ACT8* (At1g49240) was used as internal control gene for normalization. Primer information is shown in Table S5.

## DATA AVAILABILITY

The microarray data used in this article are available in NCBI GEO (National Center for Biotechnology Information Gene Expression Omnibus) and can be accessed with GSE228396.

## ACKNOWLEDGEMENT

We thank Prof. Brad Day for his generous gift of *ADF1-4Ri* lines. We also thank Prof. Richard B. Meagher for his gift of *act2/8* and *act7*, Prof. Masao Tasaka and Dr. Takehide Kato for generous gift of *fiz1*, and Prof. Kyoko Itoh and Ms. Yukiko Sugisawa in University of Tokyo for their help in performing microarray analysis.

