## Supplemental Figures for "*Arabidopsis thaliana* subclass I ACTIN DEPOLYMERIZING FACTORs regulate nuclear organization and gene expression"

**SUPPOTING INFORMATION**

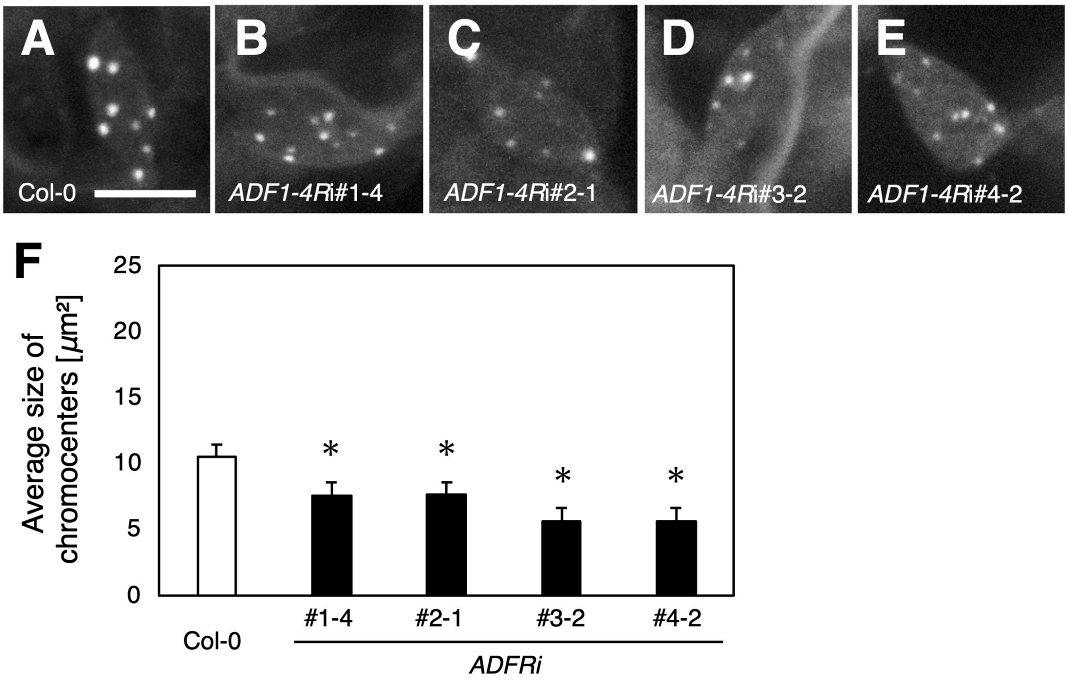


**Figure S1.** The reduction of chromocenter size was observed in all of four *ADF1-4Ri* lines. The fluorescence micrographs of DAPI-stained nuclei in epidermal cells of mature leaves in Col-0 (**A**), *ADF1-4Ri*#1-4 (**B**), *ADF1-4Ri*#2-1 (**C**), *ADF1-4Ri*#3-2 (**D**) and *ADF1-4Ri*#4-2 (**D**). Bar indicates 5 μm. (**F**) The size of chromocenters in Col-0 and four lines of *ADF1-4Ri* were quantified using fluorescence micrographs. Sixteen - 21 nuclear images were used for quantitative analysis. Comparison with Col-0 was performed by Student's *t*-test. Asterisks indicate that there is a significant difference (* $P$<0.05).

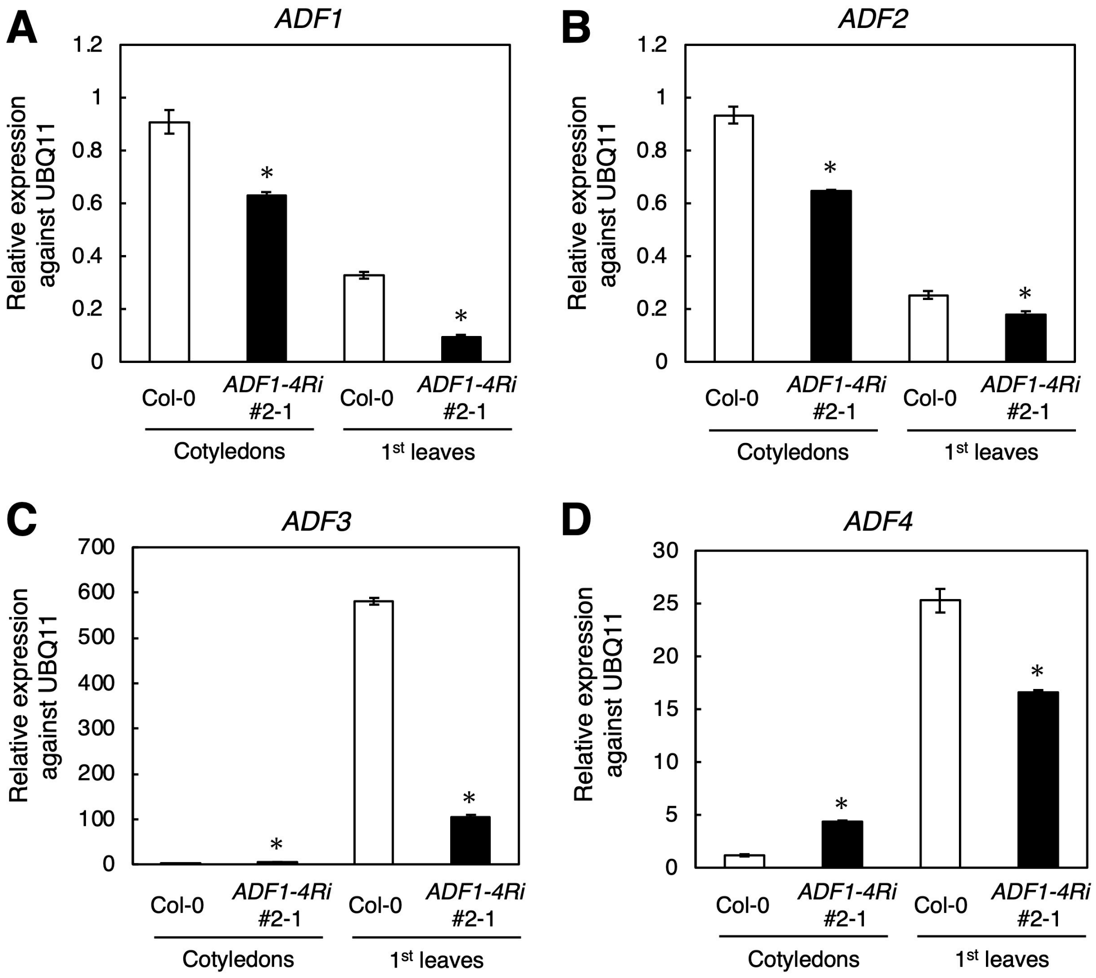


**Figure S2.** qRT-PCR using UBQ11 as a control showed suppression of expression of subclass I *ADF*s in *ADF1-4Ri* line used for Fig.2. Expression level of ADF1 (A), ADF2 (B), ADF3 (C) and ADF4 (D) in Col-0 and *ADF1-4Ri*#2-1 cotyledons and 1st leaves. Standard deviation of experimental triplicates is shown. Comparison with Col-0 was performed by Student's *t*-test. Asterisks indicate that there is a significant difference (* $P<0.05$). Representative data of biological triplicates are shown.

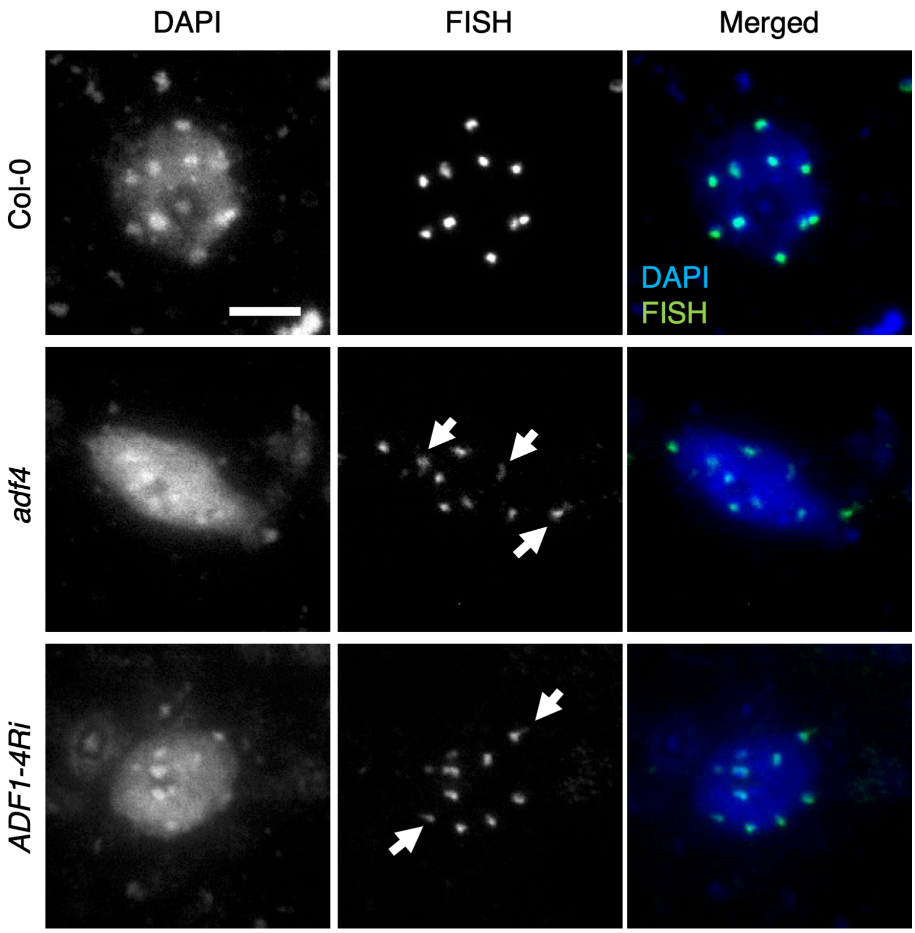


**Figure S3.** Fluorescence *in situ* hybridization analysis. The left column shows DAPI-stained nuclei, the middle column shows FISH signal, and the right column shows merged images of DAPI and FISH signal for nuclei extracted from Col-0 (top), *adf4* (middle) and *ADF1-4Ri* (bottom). Arrows indicate FISH signals that show decondensation. Bar indicates 5 µm.

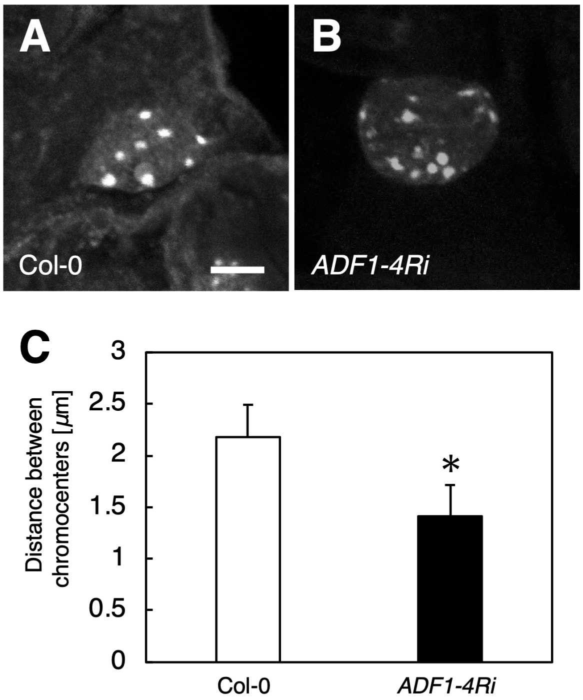


**Figure S4.** The organization of chromocenters was altered in *ADF1-4Ri*. The fluorescence micrographs of DAPI-stained nuclei in epidermal cells of mature leaves in Col-0 (**A**) and *ADF1-4Ri* (**B**). Bar indicates 5 μm. (**C**) The average of distance between two chromocenters in Col-0 and *ADF1-4Ri*. Five nuclei each were analyzed. Comparison with Col-0 was performed by Student's *t*-test. Asterisks indicate that there is a significant difference (* $P<0.05$).

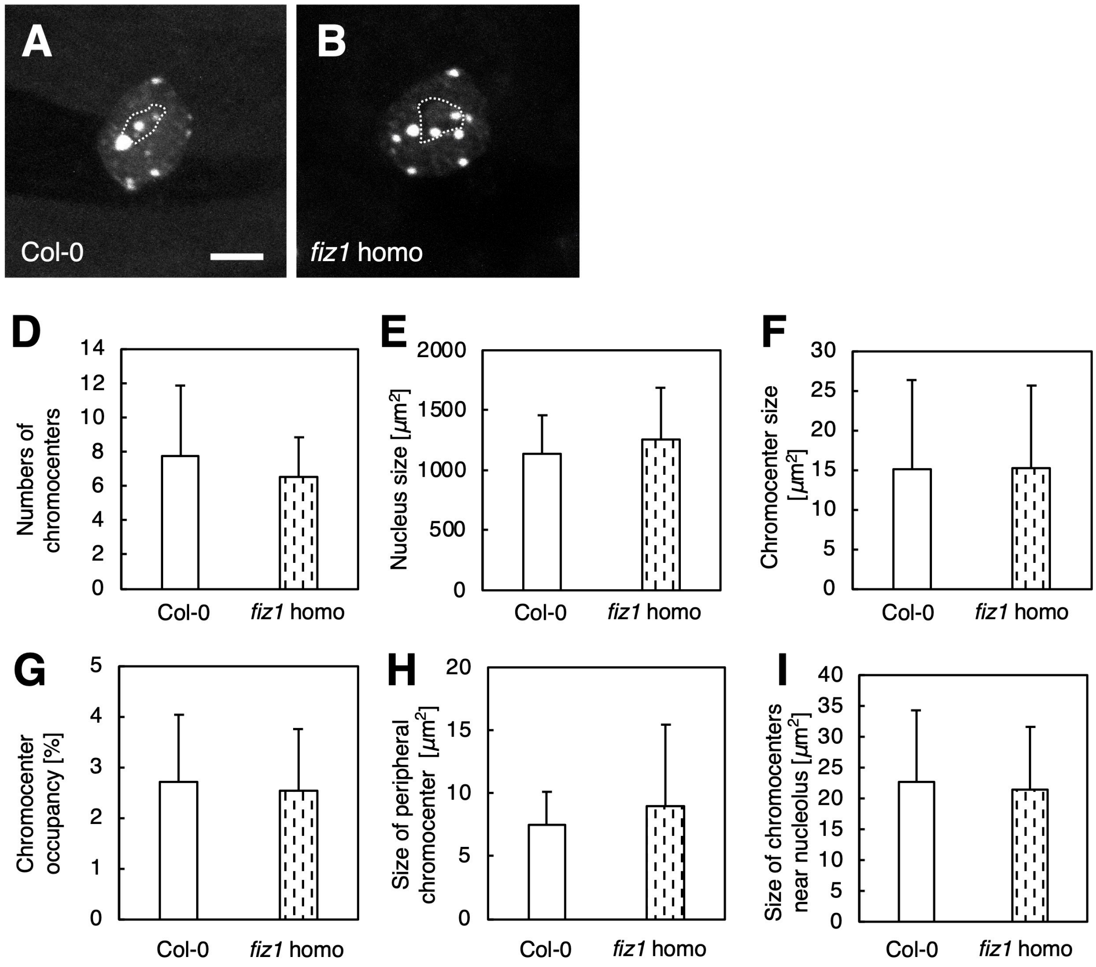


**Figure S5.** The organization of chromocenters was not altered in *fiz1* homozygous mutant. The fluorescence micrographs of DAPI-stained nuclei in epidermal cells of mature leaves in Col-0 (**A**) and *fiz1* homozygote (*fiz1* homo) (**B**). Dotted lines indicate nucleoli. Bar indicates 5 μm. (**D**) Numbers of chromocenters, (**E**) the size of nuclei, (**F**) the size of chromocenters, (**G**) the percentage of chromocenter area in nuclei, (**H**) the size of chromocenters located at the nuclear periphery, and (**I**) the size of chromocenters located near the nucleoli in Col-0 and *fiz1* homo were quantified using fluorescence micrographs. Nine (Col-0) and 10 (*fiz1*) nuclear images were used for quantitative analysis. Comparison with Col-0 was performed by Student's *t*-test and there was no statistical significance ($P$<0.05).

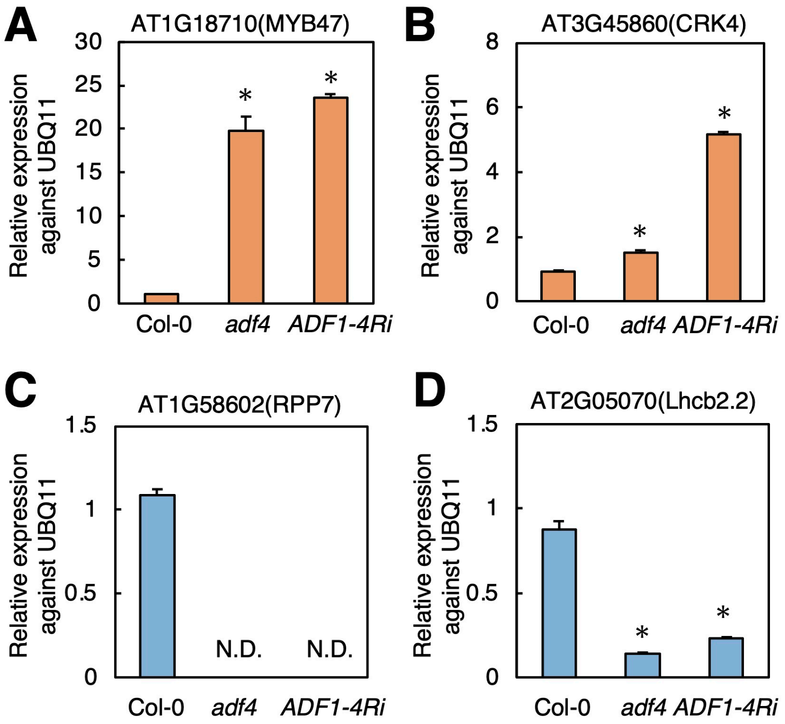


**Figure S6.** qRT-PCR using UBQ11 as a control confirmed alteration in gene expression in *adf4* and *ADF1-4Ri* shown by qRT-PCR using ACT as a control. The expression of At1g18710 (**A**) and At3g45860 (**B**) was upregulated in both *adf4* and *ADF1-4Ri*. The expression of At1g58602 (**C**) and At2g05070 (**D**) was downregulated in both *adf4* and *ADF1-4Ri*. Standard deviation of experimental triplicates is shown. Comparison with Col-0 was performed by Student's *t*-test. Asterisks indicate that there is a significant difference (* $P$<0.05). Representative data of biological triplicates are shown.

**Table S1.** Genes upregulated in both *adf4* and *ADF1-4Ri*.

**Table S2.** Genes downregulated in both *adf4* and *ADF1-4Ri*.

**Table S3.** Microarray data for *LHCB2* genes.

**Table S4.** A list of *NLR* genes that differently expressed in *adf4* and *ADF1-4Ri*.

**Table S5.** Primers for qRT-PCR.

### SUPPLEMENTAL METHODS

#### Fluorescence *in situ* hybridization

Chromosomes were prepared according to Murata and Motoyoshi (1995) and Murata *et al*. (1997) with minor modifications. The 1st/2nd leaves from two-week-old seedlings were fixed with 1:3 acetic acid:methanol fixative, and then treated with enzyme solution [(0.5% (w/v) Cellulase Onozuka RS (Yakult), 0.25% (w/v) Pectolyase Y-23 (Kyowa Kasei)] for 1 h at 37°C to yield protoplasts. After centrifugation, the protoplasts were resuspended with the above fixative, dropped onto glass slides, and then dried with heat from a gas burner.

Using the PCR DIG probe synthesis kit (Roche), the centromeric 180 bp repetitive sequences were amplified from the genomic DNA of ecotype Columbia with the following primer set: 5'-AACCTTCTTCTTGCTTCTCA-3' and 5'-GGTTAGTGTTTTGGAGTCAA-3'. The PCR products were digested with AluI and HindIII to obtain probes less than 500 bp. After ethanol precipitation, the probes were dissolved in hybrid solution (50% formamide, 20% dextran sulfate sodium, 10% 2x SSC, 5% 10 mg/ml sermon sperm DNA, 15% D.W).

Chromosomes on the slides were denatured at 80°C for 2 min in 70% (v/v) formamide in 2x SSC buffer (0.3 M NaCl, 0.03 M sodium citrate, pH 7.0). The 180 bp probes loaded onto the slides were hybridized at 80°C for 5 min and then labeled with FITC-conjugated anti-digoxigenin antibody (Roche), with the nuclei counterstained using 0.1 μg/ml DAPI (4,6-diamino-2-phenylindole).

#### Quantitative analysis of chromocenter organization

To quantify the aggregation of heterochromatin blobs, single intensity peak of each blob was picked, then searched the closest peak of most neighbor blob and measured the distance between them. We evaluate the aggregation from the median value of the distances in each nucleus. All image processing and analysis were performed by ImageJ software in combination with LPIXEL Plugins (https://lpixel.net/en/products/lpixel-imagej-plugins/).
